# SimulScan and Partial Least Squares: Visualizing swallowing through functional and dynamic imaging correlations

**DOI:** 10.1101/2025.10.29.685135

**Authors:** Bradley P. Sutton, Anthony Bosshardt, Ching-Hsuan Peng, Jiyoon Kim, Riwei Jin, Vaishnavi Krishna, William G. Pearson, Zhongming Liu, Georgia A. Malandraki

## Abstract

**Purpose:** Swallowing is a complex function involving the precise coordination of muscles, nerves, and brain areas, and can be disrupted in a variety of neurological conditions. Current imaging studies to visualize the central control of swallowing cannot examine both the biomechanics of the swallow and the brain activity associated with swallow events. An updated version of SimulScan is introduced that provides high-quality and high-speed dynamic imaging, together with fMRI acquisitions, to enable data-driven analysis of swallowing function through a partial least squares correlation (PLSC) analysis.

**Methods:** Integrating updated dynamic imaging approaches, SimulScan can achieve dynamic MRI at 23.75 frames per second and BOLD fMRI at a 1.6 s TR. Five subjects were recruited and scanned with SimulScan twice and with videofluoroscopy to compare the preliminary reliability of measuring swallowing biomechanics and the test-retest relationship in correlated functional and dynamic components of PLSC.

**Results:** High reliability of biomechanical measures of swallowing were achieved across the two SimulScan runs. In addition, these showed moderate correlation with videofluorscopy measures. Correlations between dynamic and functional imaging across runs also showed high reliability indicating that SimulScan with PLSC can extract maps of linked correlations between the brain and the oropharyngeal region.

**Conclusion:** The updated version of SimulScan with PLSC analysis enables the study of central control of swallowing, providing simultaneous biomechanical visualization of the swallow along with brain functional signals.

## Introduction

Swallowing is a complex function involving the precise coordination of oropharyngeal muscles, nerves, brainstem circuitry, and brain inputs (1). Disruptions in these elements can lead to swallowing disorders (a.k.a. dysphagia), affecting about 9.44 million adults in the US annually, often due to neurologic conditions such as stroke and Parkinson’s disease (PD) (2). Neurogenic dysphagia is linked to poor outcomes such as reduced quality of life, malnutrition, and increased respiratory infections and mortality (3–8). A perplexing clinical challenge is that neurogenic dysphagia is highly heterogeneous even within a disease (9,10), may result from involvement of many neural mechanisms (11,12), and remains relatively poorly understood (9,13). This heterogeneity and limited understanding hinder the development of effective, physiology-based swallowing treatments and inhibit patient outcomes.

To start addressing this gap and better understand the central control of human swallowing, over the last few decades studies have used a variety of neuroimaging methods. Among them, the use of task-based functional MRI (fMRI) has been prominent. The vast majority of these studies have investigated healthy adults and have showcased an extensive bilateral network of cortical and subcortical areas involved in this critical biological act, including the lateral primary somatosensory and motor cortex, the premotor area, the inferior frontal gyrus, as well as areas in the basal ganglia, the anterior cingulate, insula, cerebellum, and components of the brainstem (for a meta-analysis, see (14)). Task-based fMRI and resting state functional connectivity MRI (fc-MRI) have also been increasingly used to examine the effects of neural disruptions, such as those caused by stroke and PD, on the larger sensorimotor network (15–18), as well as to study swallowing neurophysiology in patient populations and brain plasticity related to treatments in patients with dysphagia (19–22).

In studies using task-based fMRI to examine swallowing control in stroke patients with dysphagia, results have been mixed, showing both hyperactivation and hypoactivation of key cortical areas compared to activity in healthy controls (e.g., (23–26)). Pre-post treatment/recovery studies have similarly provided preliminary data on changes in resting-state or task-based activity in specific brain regions that seem to be, at least partly, related to stroke size and severity (22,27–29). Although these studies have offered valuable initial information into central adaptations in neurogenic dysphagia, key challenges in applying this methodology have limited the number of participants and the overall number of studies in this field. These challenges primarily relate to the motion artifacts that are often introduced when patients swallow stimuli in the magnet, as well as the inability of patients with dysphagia to safely swallow under artificial conditions – e.g., at exact times and with specific stimuli, as required for accurate fMRI imaging and analysis.

Furthermore, the task-correlated motion and magnetic susceptibility changes during swallowing is a significant challenge for fMRI studies (30). This motion will show up as false fMRI activation, usually occurring around the periphery of structures where signal change will be highest from the artifact. Several strategies have been used to try to avoid motion in swallowing studies. In (31), a behavioral interleaved gradient method (32) was used to place the swallowing motion in a quiet period of the functional MRI acquisition, limiting the impact of motion on the data. This, however, made the data acquisition less efficient with consistently longer TRs required to allow time for motion. Other approaches to minimize motion include adding extra head padding to minimize movement, monitoring motion during the scan to exclude data with excessive motion, and tracking precise timing of the swallow to attempt to separate motion and neural signals. In addition, there are navigator acquisitions that measure some physiological signal fluctuations with an additional acquisition to correct each functional image (33). Despite these approaches, there is still a need for motion-robust acquisitions that can capture swallowing motions and separate neural activations from motion-induced signal fluctuations.

In order to understand the impact of central control on swallowing, there is a need to visualize the specific biomechanical deficits in swallowing function, as they provide critical information for diagnosis and treatment planning. In most prior fMRI studies, swallowing was monitored roughly through the use of monitoring devices such as surface electrodes or pneumographic belts in the neck area (31,34,35). These methods can only indicate whether a swallow occurred without providing any diagnostic information on swallowing physiology or biomechanics. Further, they likely introduce sensory feedback, potentially impacting typical motor and sensory swallow function (36). Current standard of practice for the diagnosis of dysphagia involves traditional imaging approaches like videofluoroscopy (real-time x-ray) and endoscopy, with both methods offering real-time anatomical and physiological imaging of the swallow. They further require the use of scoring systems and scales (37,38) that are based on the subjective visuo-perceptual interpretation of the videos/images by clinicians. At this time, neither of these approaches enables the examination of the associated brain activity during the swallow to determine the impact of central control on a functional or dysfunctional swallow. Instead, to date, clinicians and researchers have been limited in acquiring brain activation information through MRI/fMRI or medical reports and swallowing imaging information from x-ray/endoscopy separately and deriving relative correlations between the two (e.g., (11,12,21,26,39)). These mostly correlational attempts have left us with multiple unanswered questions about the specifics of the central swallowing control. For example, we still lack a complete understanding of the specific neurophysiological factors that drive swallowing events and symptoms, how these factors interact, and what neuromuscular and neurophysiological adaptations occur in different patient groups. Recognizing these factors and adaptations is considered a crucial step in our pursuit of creating more effective and personalized treatments for neurogenic dysphagia (13,40,41).

The SimulScan sequence was introduced in order to address the need for motion-tolerant fMRI of central control of swallows while simultaneously evaluating the biomechanics of those swallows and eliminating the need for a stimulus swallow in the supine position (36). The sequence interleaved BOLD-based fMRI with a dynamic imaging sequence to visualize motion associated with incidental swallows during a long, covert, fMRI study (36). Although the sequence was able to achieve acquisition of fMRI and dynamic MRI, the dynamic data only had sufficient quality to visualize the timing onset of swallows for the covert swallowing task. Since publication of the SimulScan technique, new advances in low rank and Partial Separability (PS) models have enabled drastic increases in imaging speed, spatial resolution and image quality (42). These techniques have been used to create dynamic MR imaging studies that provide high-quality, high temporal resolution images of speech and swallowing with image fidelity similar to structural scanning (43–47).

Given the potential for SimulScan to provide insights into neural components of swallowing changes with age and disease, we present an updated acquisition that enables high quality dynamic MRI enabling biomechanical analysis while simultaneously acquiring BOLD-based fMRI. In addition, leveraging the richness of the dynamic and functional data, we demonstrate a data-driven approach to extract correlated brain function and dynamics from the dataset. To demonstrate its reliability, we perform a test-retest reliability analysis from a set of healthy young participants. In addition, we demonstrate preliminary validation in tracking dynamic anatomical landmarks through computational analysis of swallowing mechanics (CASM) (48,49) using dynamic MRI compared to the standard of practice tool (videofluoroscopy) in the same participants.

## Methods

In this work, an updated SimulScan imaging acquisition and reconstruction method are implemented that leverage recent advances in dynamic imaging, drastically improving the image quality of the dynamic MRI portion of SimulScan. In addition, an analysis approach leveraging Partial Least Squares Correlation (PLSC) is used to analyze the dynamic and functional neuroimaging data together in a data-driven manner. We demonstrate that the new dynamic images from SimulScan can be reliably tracked through a Computational Analysis of Swallowing Mechanics (CASM) approach and are comparable to swallows acquired using videofluoroscopy in the same subjects. Finally, to demonstrate reliability of the SimulScan method, we perform a test-retest experiment and analysis.

### Updated Acquisition

The SimulScan sequence (pulse sequence diagram in Fig. 1) interleaves a mid-sagittal dynamic imaging sequence with a BOLD-based fMRI acquisition collected with oblique axial slices in the brain. An overall TR (42.1 ms) includes a block for a sagittal dynamic acquisition (achieving 23.75 frames per second (fps)) and a block for a single slice of fMRI data (BOLD TR of 1.6 s for 38 slices). For the dynamic imaging portion of the sequence (Fig. 1B), leveraging advances in PS-model speech imaging (45,46), we combine a spiral-in navigator that is repeated each dynamic block, followed by a spiral-out shot that rotates in each acquisition. This is immediately followed by a spiral-in BOLD-based fMRI acquisition with a TE of 25 ms acquiring 1 shot for a single 2D slice.

**Figure 1:**
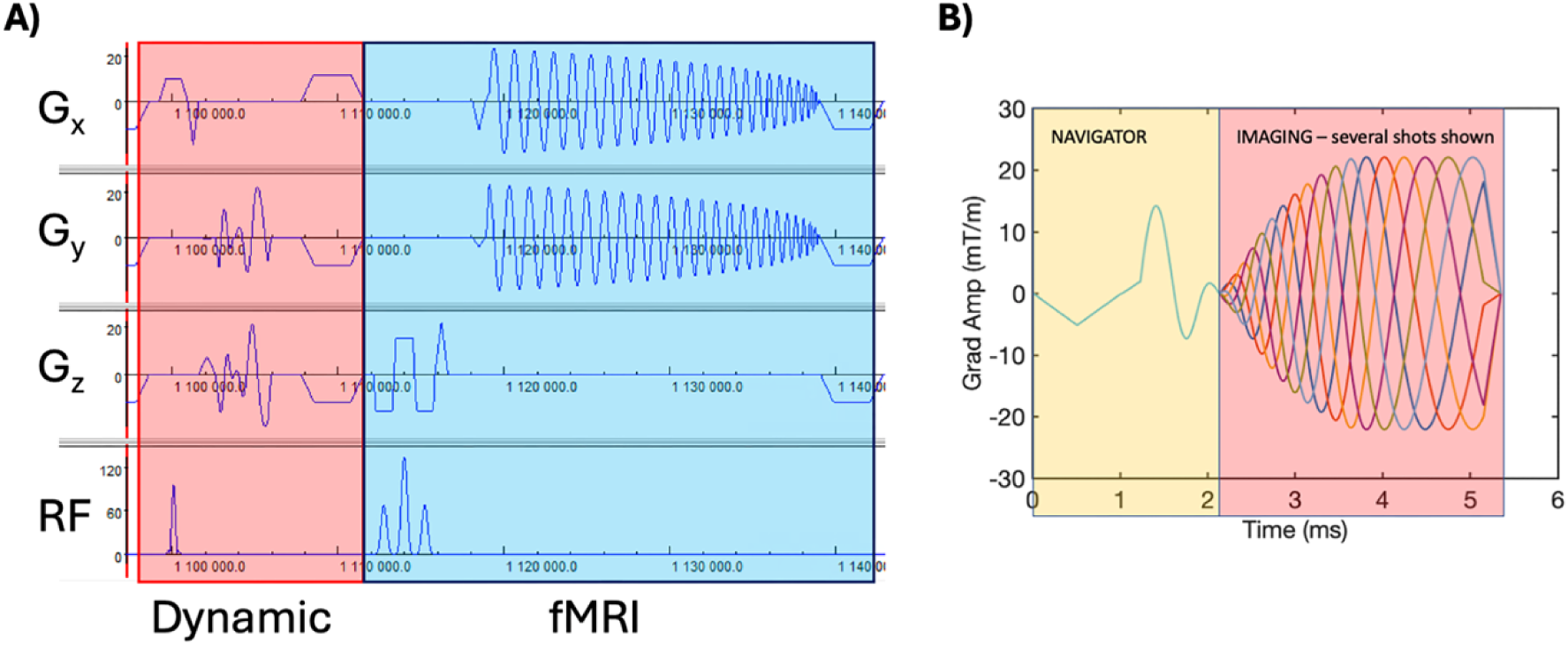
**A)** Pulse sequence diagram for the SimulScan sequence. Red-shaded portion is the sagittal dynamic imaging and blue shaded region is the 2D axial fMRI acquisition with TE of 25 ms. The fMRI acquisition uses water excitation from a binomial pulse and a spiral-in acquisition. **B)** The dynamic imaging module (expanded to clearly display the two components) consists of a spiral-in navigator that is repeated each time followed by a single shot of a spiral-out acquisition. Several shots are shown overlaid.

In our imaging protocol, the SimulScan sequence acquired a dynamic image at 23.75 frames per second for a single mid-sagittal 10-mm thick slice. The spiral-in temporal navigator was 1-shot of a 40-shot acquisition for a 30 cm FOV with a 3 mm spatial resolution, designed with a maximum gradient amplitude and slew of 23 mT/m and 120 T/m/s. The spiral-out dynamic imaging data was a multishot spiral acquisition designed for a 30 cm FOV, 200 matrix size (but reconstructed at 225 for 1.3 mm in-plane resolution), 80-shot spiral with the same maximum gradient and slew rate. For the dynamic imaging sequence, we use an RF pulse with a flip angle of 10° and use both gradient and RF spoiling. Simultaneously, fMRI scans were taken for 38 slices, 3 mm thick, with in-plane resolution 2.5x2.5 mm and matrix size of 96x96, acquired with a spiral-in acquisition that was 1 shot of a 2-shot acquisition, parallel imaging acceleration factor of 2, and a 90° flip angle binomial water excitation RF pulse. The TR for the fMRI portion of the sequence is 1.6 s.

From this acquisition, traditional 2D fMRI slices are reconstructed using a magnetic field inhomogeneity (field map measured in separate acquisition) corrected image reconstruction (50). Dynamic data are reconstructed using the PS-model reconstruction framework (45,46). Briefly, the spiral-in navigator data are formed into a k-space by time-point matrix, and the top 40 right singular vectors are taken as the temporal basis. Then the spiral-out dynamic imaging data are used to fit the spatial basis functions to the corresponding temporal basis through a least-squares framework. The spatial basis functions are combined with the temporal basis functions to create a full timeseries of images.

### Partial Least Squares

With the updated SimulScan sequence, we can achieve a rich set of dynamic imaging data and the associated brain activity underlying the activity. To analyze central control of swallowing, we need to extract the brain activity related to the swallows in the data, but we also have potential subject bulk motion contamination in the fMRI data during the swallow. This requires a data driven approach to find dynamic relationships between the swallowing and fMRI data.

Partial Least Squares Correlation (PLSC) has been used previously to analyze relationships between dynamic behavior and functional activations in fMRI data (51,52). In this method, an SVD is used to extract latent variables of correlated components from the dynamic imaging and functional brain activation from a correlation matrix between the behavioral and fMRI data. Let the fMRI signals be stored in a matrix ***X***, which is a matrix that is *N_t_* (number of total fMRI TR’s in the SimulScan run) by *N_x_* (the number of voxels in the 3D brain images). And the dynamic imaging data will be stored in ***Y***, a matrix that is *N_t_*by *N_y_* (the number of voxels in the 2D mid-sagittal dynamic imaging slice (or a region of interest in that slice)). We note that *N_t_*must be the same between the dynamic imaging series and the fMRI data in our current implementation of PLSC. To accomplish this, we let the dynamic information in ***Y*** be the temporal standard deviation in each voxel’s magnitude across the fMRI TR, which is comprised of 38 dynamic images. Further, we incorporated the hemodynamic delay in the dynamic time series to achieve aligned dynamics and brain signals. The temporal standard deviation in the new TR windows was the dynamic signals comprising the matrix ***Y***.

We follow the formulation for the behavior approach for PLSC (52), which is summarized here. See Fig. 2 for an overview of the method. The correlation between the dynamic imaging data and the fMRI signals is formed by taking ***R = Y^T^X***. Taking the SVD of the correlation matrix, **R**, results in a set of singular vectors, or saliences, ***U*** and ***V***. We choose a particular rank *L* to examine only the strongest correlated components. These singular vectors are the correlated spatial maps related to brain function (or functional activation maps for the correlated component) in the right singular vectors of R or the columns of ***V***, a *N_x_* by *L* matrix. And the associated dynamic heat maps in the left singular vectors of ***R***, or the matrix ***U***, a *N_y_* by *L* matrix.

**Figure 2:**
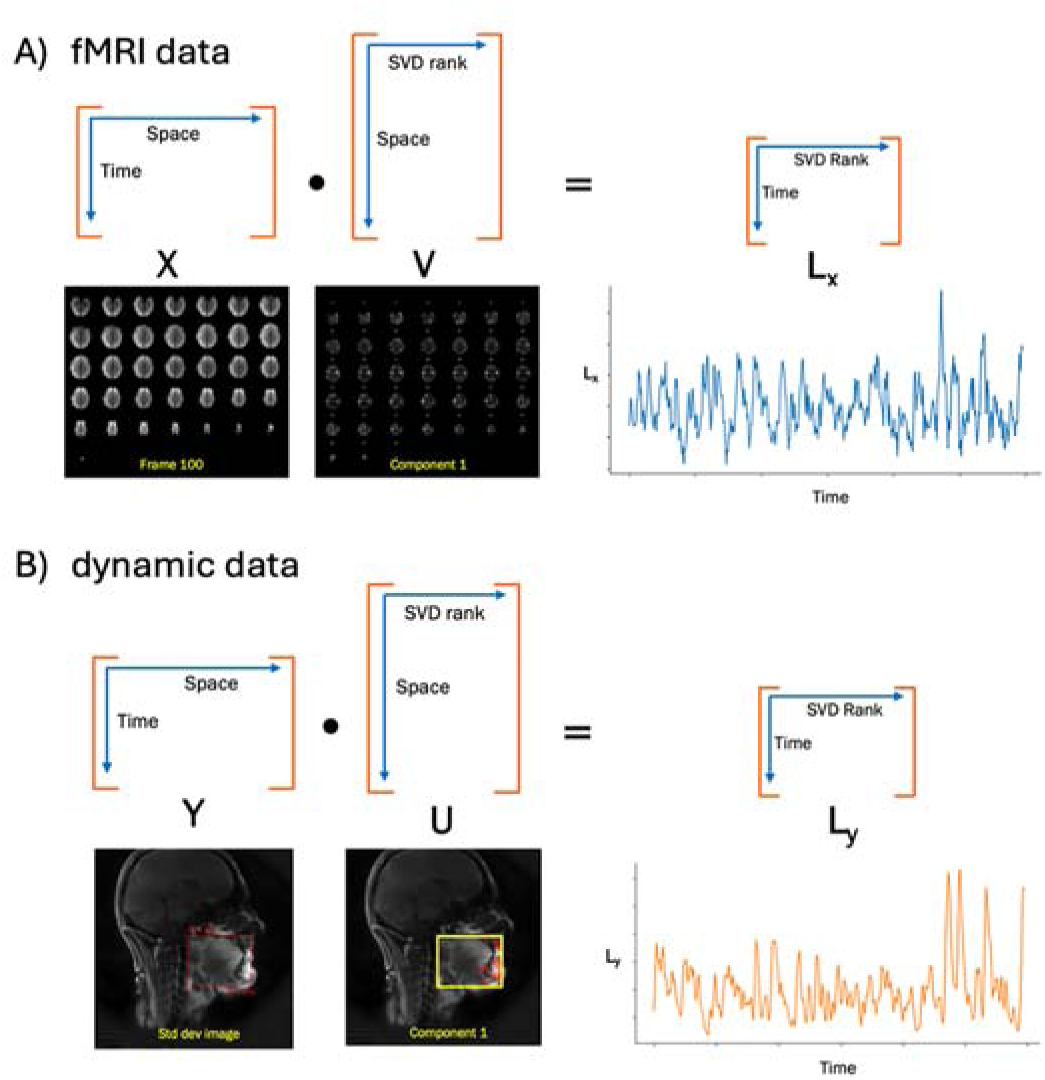
Formulation for the behavioral partial least squares correlation (PLSC) approach, where the correlation, R, is determined from the fMRI data, X, and dynamic data, Y, as R=Y’X. **A)** Layout of the fMRI data in matrix X, and the matrix V extracted maps from the SVD of the correlation matrix. One component of V is shown. This is used to form the latent variable time series L_X_. **B)** Layout of the dynamic imaging data, Y, along with the spatial maps, U, extracted from the SVD of the correlation matrix and its latent variable time series L_Y_.

We can extract temporal waveforms showing the dynamics of correlated components in the dynamic and fMRI data by forming the latent variable timeseries as follows:

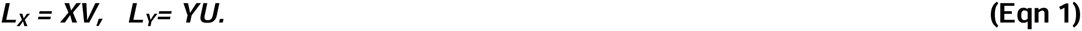

These latent variables, ***L_X_*** and ***L_Y_***, are *N_t_* by *L*, giving the time series for each correlated component. Note that (Eqn 1) represents applying each individual component’s spatial map (activation map or heat map) to the respective time series data to extract a weighted time series of that map’s activity.

Due to the large size of the data, custom Python libraries were written to provide the PLSC analysis and other processing. This script is available on GitHub (provided at time of acceptance). A custom Python decorator function was written to report RAM usage for arbitrary function calls, providing immediate feedback on the memory efficiency of computationally intensive operations. The MATLAB implementation of SVD of ***R*** requires loading the entire input matrix into memory and peak RAM usage during computation approached 10 times the size of the input, creating memory issues on a 256 GB RAM workstation. In contrast, the Dask implementation in python computes a randomly compressed rank-*L* thin SVD, yielding only the user-specified leading *L* singular values (53). In practice, the Dask implementation consumes approximately 20 times less RAM than the MATLAB implementation, enabling execution on a wide range of consumer hardware.

### Evaluation of SimulScan

In order to evaluate the ability of the SimulScan sequence to capture dynamic motions associated with swallowing and to assess the reliability of fMRI components associated with swallowing, SimulScan data was acquired on 5 healthy young subjects (mean age = 25 ± 3.46 years; 3 female) in accordance with a protocol approved by the institutional review board at Purdue University (IRB# 2023-714). All participants were healthy, in that they had no cognition deficits based on a cognition screening tool, Montreal Cognitive Assessment (MoCA) (54), and no swallowing concerns based on a swallowing screening tool, Eating Assessment Tool (EAT-10) (55). Participants were excluded if they had a history of head and neck cancer or neurological disease, or any condition which would contraindicate MRI. Data was acquired on a Siemens 3T Prisma MRI scanner with a 64-channel head coil at the Purdue Life Science MRI facility in West Lafayette, Indiana, using the SimulScan sequence along with T_1_ and T_2_ structural scans of the brain and oropharyngeal region.

Similar to our earlier work, a simple saliva swallowing paradigm was used while subjects watched a video of their choice. 440 BOLD TR’s were acquired per self-paced swallowing SimulScan run, and two runs were acquired for each subject to allow test-retest examination of extracted correlated components. For one subject, we acquired 600 BOLD TR’s per run (Subject 001), but shortened to 440 for all other runs. For each run, subjects were instructed to think of something tasty (to elicit saliva production), and swallow their saliva as frequently as comfortable throughout the acquisition, with no target number of swallows given, resulting in a variable number of swallows per run. Across all subjects, there was an average of 29 spontaneous swallows per scan. Subjects were padded into the head coil to minimize head motion and were instructed to keep their head still. Swallows were saliva swallows, no stimuli were given to the subject. fMRI data from SimulScan were processed in a typical preprocessing pipeline in FSL (6.0.5.1, (56)), including: motion correction using MCFLIRT (57); spatial smoothing using a Gaussian kernel of FWHM 5 mm; and highpass temporal filtering (Gaussian-weighted least-squares straight line fitting, with sigma=50.0s).

### Dynamic Tracking Comparison using CASM

Subjects who participated in the MRI experiment also performed swallows under videofluoroscopy (VFSS). VFSS was acquired using a videofluoroscopic C-arm system (OEC 9800 Plus Digital Mobile 12 in. GE), which records images at 30 frames/second at the highest resolution (30 pulses/second). Subjects were placed in the supine position and performed 3 saliva swallows, however, these swallows were cued, which is necessary to reduce ionizing radiation exposure.

For the preliminary validation included herein, and given the large number of swallows completed during the SimulScan scans, we randomly selected and analyzed two swallows from the VFSS and two swallows from each SimulScan run for comparison. For both the dynamic MRI images and the VFSS swallows, a semiautomated MATLAB tracking tool (58) was used to annotate 10 coordinates of anatomical landmarks mapping the functional anatomy of pharyngeal swallowing mechanics. Coordinates included five soft tissue landmarks (upper esophageal sphincter [UES], inferior edge of posterior vocal folds, inferior edge of anterior vocal folds, anterior inferior edge of hyoid bone [hyoid bone], and base of tongue/valleculae]) and five hard tissue landmarks (inferior genial tubercle of mandible [mandible], intersection of posterior edge of hard palate and pterygoid plate of sphenoid bone [hard palate], anterior tubercle of first cervical vertebra [C1], anterior and inferior edge of the second cervical vertebra [C2], and anterior and inferior edge of fourth cervical vertebra [C4]) (48). These landmarks are manually identified at swallow onset and then semi-automatically tracked frame-by-frame to to record coordinate data for multivariate morphometric analysis. Fig. 3 displays the anatomical landmarks annotated on a frame of a VFSS and SimulScan, respectively.

**Figure 3:**
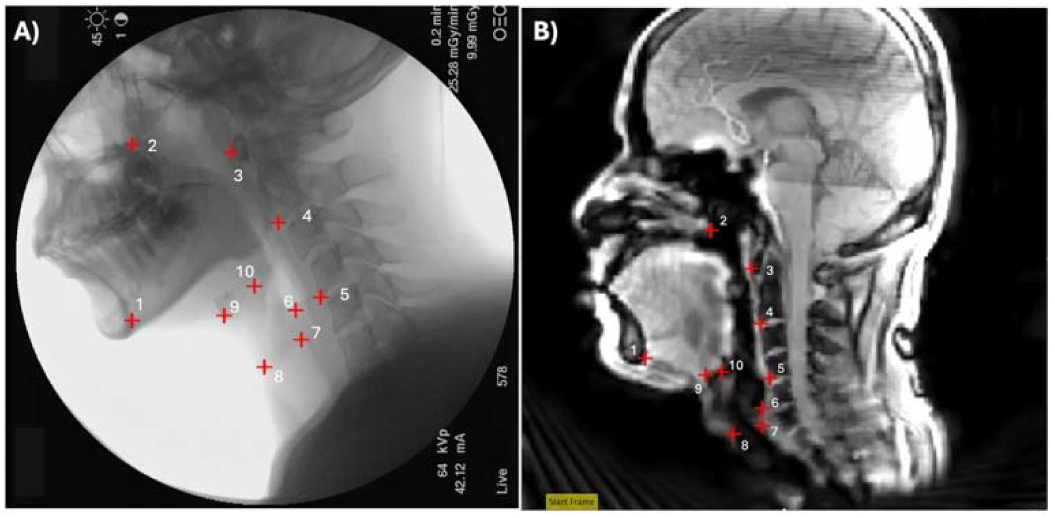
The landmarks used in Computational Analysis of Swallowing Mechanics (CASM). **A)** The 10 anatomical landmarks tracked via CASM in VFSS. **B)** The 10 anatomical landmarks tracked via CASM in dynamic MRI.

All CASM raters met > 0.95 agreement with the method developer (author WP) on a training set prior to data analysis. Swallow videos were first trimmed to predefined start and end frames prior to CASM analysis (48,59). Because this study is the first application of CASM to saliva swallows a new standardized trimming protocol was developed. Start and end frames for each image modality were determined differently to account for differences in temporal resolution and modality-specific fidelity in visualizing anatomical structures. For VFSS, the first frame was defined as 3 frames before the velum starts elevating, and the end frame was defined as 10 frames after the hyoid bone starts moving posteriorly and inferiorly from its highest position. The start frame for SimulScan was defined as 7 frames before the first frame showing the onset of tongue base to posterior pharyngeal wall contact, and the end frame was defined as 3 frames after the last frame of tongue base to pharyngeal wall contact.

Matrix correlation analysis using Morpho J (60) was conducted to examine the correlations between the coordinates annotated in SimulScan and VFSS, as well as between the two SimulScan runs. Discriminant functional analysis (DFA) was used to further explore the landmarks that drive overall pharyngeal shape differences. DFA output was then imported into the MATLAB tracker tool and spatially adjusted using the C1, C2 and C4 vertebrae as anchor points to reorient the shape relative to fixed skeletal structures, enabling anatomically accurate visualization and interpretation (48).

### Reproducibility analysis

To examine the reliability in extracting components of central control of swallowing, we leveraged the two runs of SimulScan in a test-retest analysis. In each run, the number and timing of swallows are different, however, if we are extracting functional regions involved in swallowing control, then we would expect that the same brain regions are correlated with the same regions of dynamic motion in the second scan compared to the first. For our test-retest analysis, we take the functional brain maps, ***V***, from run 1 and apply it to the functional brain imaging data, ***X***, in run 2. By doing this, we extract a cross-run latent variable time series ***L_X_*** from run 2. We also take the dynamic heat map, ***U***, from run 1 and apply it to the dynamic imaging data, ***Y***, from run 2, forming a cross-run latent variable time series ***L_Y_*** from run 2. If we have extracted correlated components, then the two latent variable time series we extracted with those masks should still be correlated. We examine this cross-run latent variable correlation compared to within-run correlation for all 4 pairings that are possible: two within run (data from run 1, maps from run 1; data from run 2, maps from run 2) and two across run (data from run 1, maps from run 2; data from run 2, maps from run 1). This is shown schematically in Fig. 4. We use Pearson correlation between pairs of timeseries.

**Figure 4:**
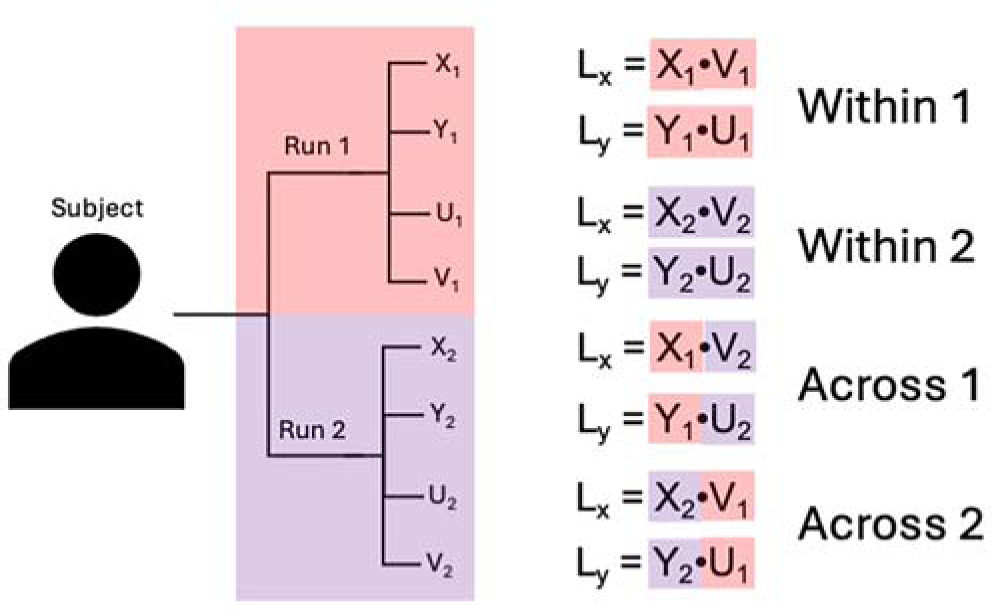
Schematic of test-retest comparison of Simulscan dynamic and functional imaging data. We form within run correlations, using maps and data from the same run, shown in the first 2 rows. We also do cross-run correlations using maps from 1 run and data from the other, shown in the bottom 2 rows.

## Results

Fig. 5 shows example images from the reconstruction of the SimulScan acquisition on a subject. In an 11 min 44 sec scan, 440 functional volumes are acquired and 16,720 dynamic frames of the mid-sagittal slice. The functional images show a dark line down the middle where the dynamic mid-sagittal image is acquired. Subjects were swallowing at their own pace and a time series of 14 images (589 ms) shows one of the swallows that occurred during the run (Fig. 5C).

**Figure 5:**
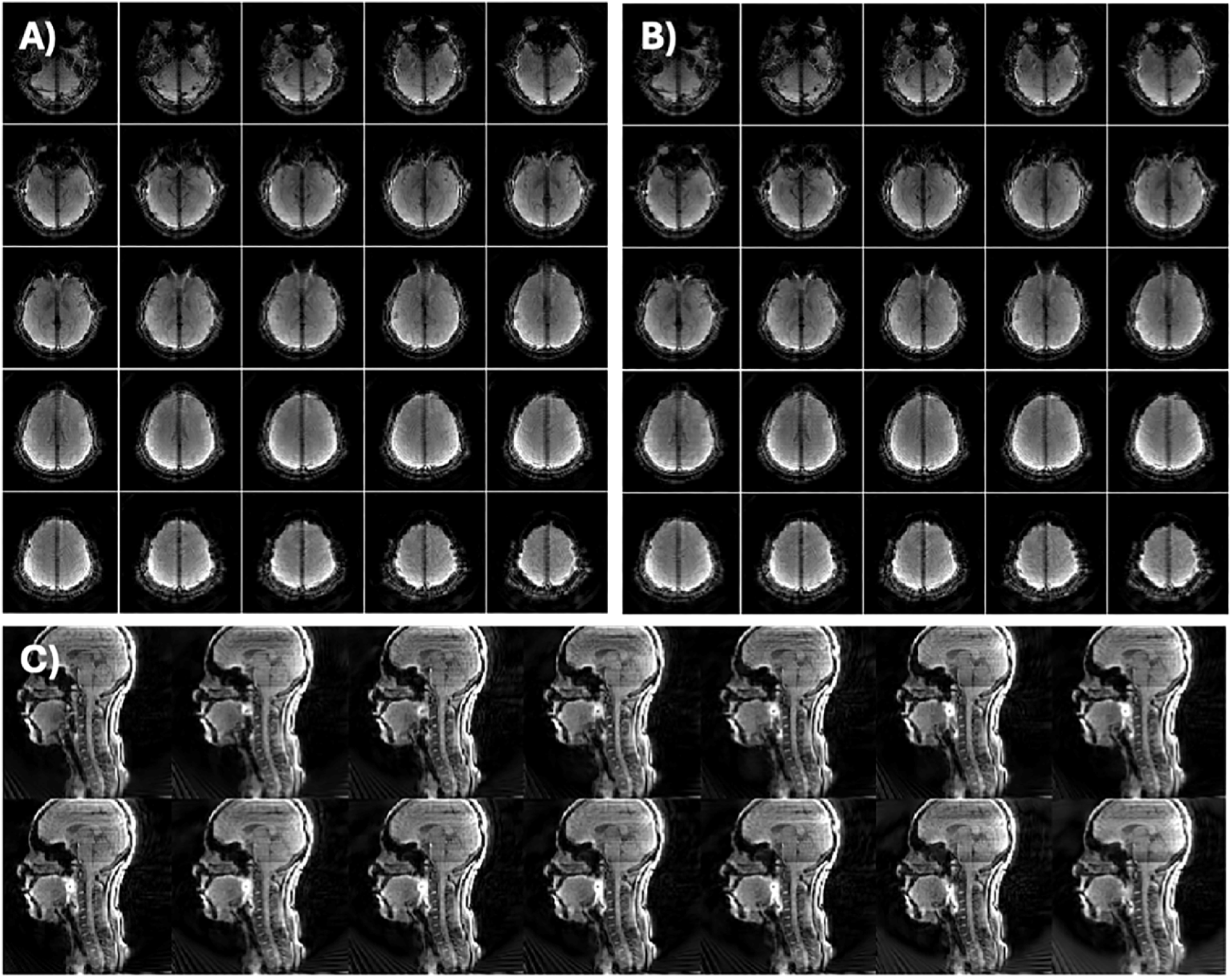
Example output from SimulScan. **A-B)** Middle 25 slices shown from the BOLD fMRI scan at two time points during the acquisition, randomly selected and 1.5 minutes apart. **C)** 14 dynamic frames (589 ms) during a swallow in the same SimulScan run.

PLSC analysis was performed on all runs from the study, extracting the top 10 correlated components. Fig. 6 shows several components (top 3) from two subjects that were extracted by PLSC from within Run 1. The activation or heat maps for the fMRI and dynamic data are shown with thresholding the largest magnitude (positive and negative) 10% of correlation intensities. Coefficient maps were each normalized by their maximum value before display. These masks are visualized as the colored overlays shown in Fig. 6 for both brain function and dynamic image slices, where 4 fMRI slices were extracted for ease of visualizaiton. This shows significant correlations between swallowing events and brain activation areas. Note that the brain functional map for component 1 in Subject 1 indicates a correlation between movement of the superior tongue surface and the velum correlated with primary motor cortex activation in the brain. Component 2 reflects the pharyngeal portion of the swallow and Component 3 shows changes that may be related more to bulk motion.

**Figure 6:**
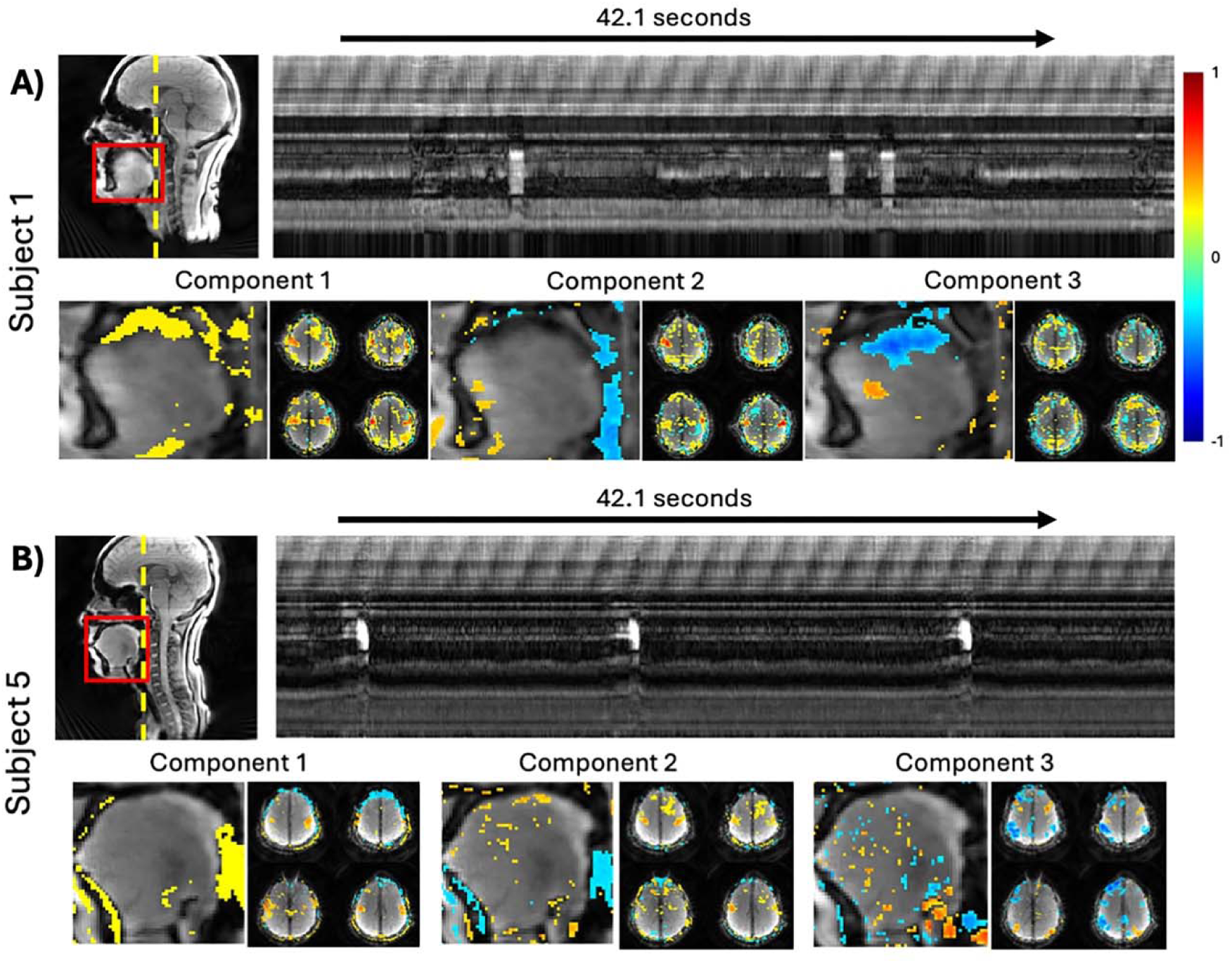
PLSC components for one run in two subjects. **A)** Subject 1 showing dynamic reference image and strip plot for 42 seconds chosen to show several swallows. Bottom row shows the scaled dynamic and functional maps from the first three components of the PLSC analysis, showing the highest magnitude 10% of coefficients in each mask. **B)** Subject 5, their strip plot chosen to show multiple swallows in a 42 second window and the fMRI and scaled dynamic MRI component maps from the first three components.

In order to assess if the improved dynamic imaging from SimulScan is sufficient to track 10 key anatomical landmarks of the swallow biomechanics, CASM was applied to both the dynamic MRI and the VFSS data. A strong positive correlation was observed between the two SimulScan runs (*r* = 0.891; *p* < 0.0001; Fig. 7A). Correlations between tracking measures for VFSS and dynamic MRI in this sample demonstrated a moderate positive correlation (*r* = 0.686, *p* < 0.0001; Fig. 7B). We note that these analyses reflect different swallows as the VFSS and dynamic MRI are performed separately, but they were performed on the same day, with VFSS preceding the MRI scan for each subject. To further explore the landmarks’ contributions to these results, discriminant functional analysis was used to compare differences in biomechanics reflected in these two modalities (Fig. 8). In Fig. 8, eigenvectors at each landmark reflect the direction and magnitude of mean variance of each coordinate. Larger magnitudes were observed with the five hard tissue landmarks, including mandible, hard palate, C1, C2, C4, and one soft tissue landmark, UES. The results indicate that these landmarks are driving the shape differences between the two modalities.

**Figure 7:**
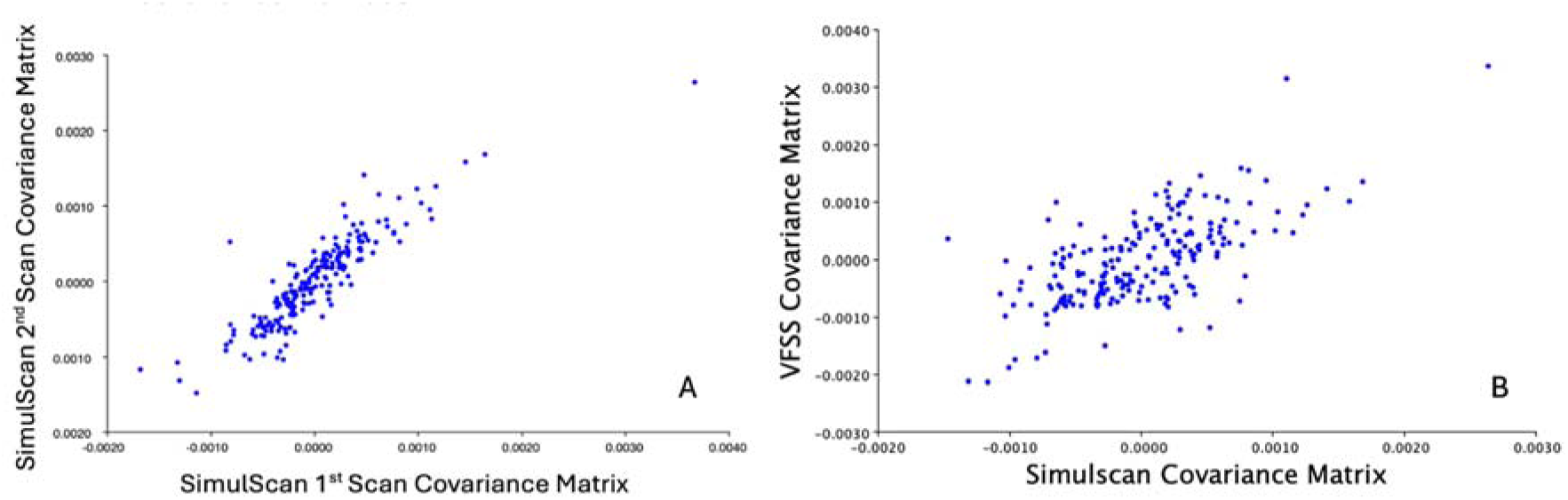
**A)** Matrix correlation between first run of SimulScan and second run of SimulScan Procrustes covariance matrices. **B)** Matrix correlation between SimulScan and VFSS Procrustes covariance matrices.

**Figure 8:**
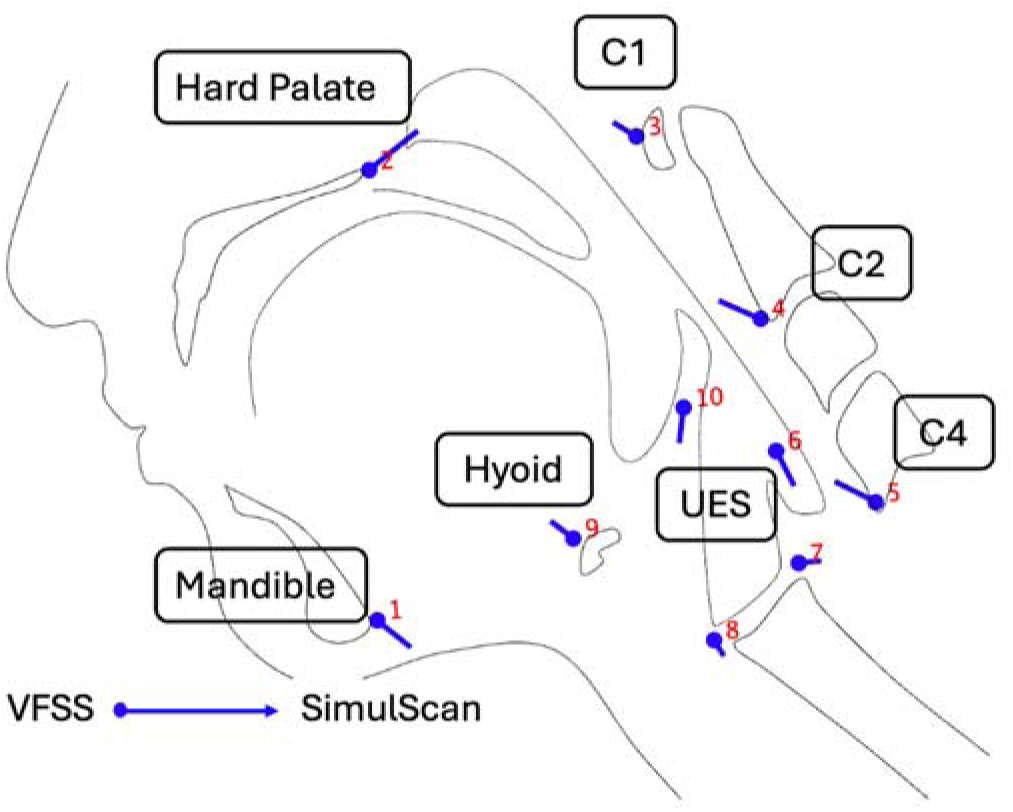
Eigenvectors from a discriminant function analysis illustrate greater differences in hard structures (mandible, hard palate, C1, C2, C4) and soft structures (hyoid bone, UES) when compared across imaging modalities.

The component maps extract the correlation regions between brain function and dynamic motions uring the swallow. To assess reliability of the correlations between the functional brain maps and the dynamic heat maps extracted by PLSC, we examined correlations between the resulting latent variable timeseries for maps used both within and across runs. The correlations for all five subjects are given in Table 1 for the first 3 components, where all correlations are significant at p<0.001 except for a few that are marked. Nearly all the cross-pairings demonstrate significant correlations (25/30) despite the functional maps and heat maps coming from different runs than the actual dynamic and functional imaging data with different timings of the swallows.

**Table 1:** Examination of use of correlated maps within and across runs. The table shows the correlation coefficient between the latent variable time course for the functional data, L_X_, and the dynamic imaging data, L_Y_, for cases in which the maps (U, V) came from the same run or a different run as the data of the SimulScan sequence. All correlations are significant at p<0.001, except those marked. x indicates not-significant, * indicates significant at p<0.01 but not p<0.001.

| Subject<br>(Components 1-3) | | $X_1U_1 - Y_1V_1$<br>Within Run 1 | $X_2U_2 - Y_2V_2$<br>Within Run 2 | $X_1U_2 - Y_1V_2$<br>Cross Run | $X_2U_1 - Y_2V_1$<br>Cross Run |
| --- | --- | --- | --- | --- | --- |
| Subject 1 | C1 | 0.6665** | 0.4909 | 0.1592 | 0.3289 |
|  | C2 | 0.5745** | 0.5680 | 0.4858 | 0.5564 |
|  | C3 | 0.4741 | 0.4557 | 0.1663 | 0.2006 |
| Subject 2 | C1 | 0.6823 | 0.6857 | 0.4049 | 0.3170 |
|  | C2 | 0.4975 | 0.4823 | 0.1079 * | 0.2617 |
|  | C3 | 0.6288 | 0.4447 | 0.1908 | -0.0230 <sup>x</sup> |
| Subject 3 | C1 | 0.5974 | 0.5020 | 0.1430 | 0.3365 |
|  | C2 | 0.6198 | 0.4745 | 0.1790 | -0.0379 <sup>x</sup> |
|  | C3 | 0.3631 | 0.5791 | -0.0329 <sup>x</sup> | 0.1941 |
| Subject 4 | C1 | 0.5359 | 0.3955 | 0.1521 | 0.3443 |
|  | C2 | 0.3982 | 0.5013 | 0.2776 | 0.2154 |
|  | C3 | 0.3764 | 0.3556 | 0.0816 <sup>x</sup> | 0.1332 * |
| Subject 5 | C1 | 0.6574 | 0.5359 | 0.4070 | 0.3034 |
|  | C2 | 0.4309 | 0.4162 | 0.4606 | 0.4377 |
|  | C3 | 0.4215 | 0.5226 | -0.0176 <sup>x</sup> | 0.1270 * |

In order to further demonstrate the motion robustness of our acquisition and analysis approach, we reanalyzed the data from one subject with a traditional fMRI approach. Using the dynamic time series, we placed a control voxel in the posterior oropharyngeal region (see Fig. 9) and extracted the time series. Since this region has tissue in it during a swallow, a high signal in this voxel’s time series indicates a swallow is occurring. We used this timing to determine task timing for a GLM analysis. With this analysis, we obtained the standard activation map shown in Fig. 9C, which has obvious motion corruption. For comparison, we also include several functional brain map components from the same run demonstrating that SimulScan with PLSC analysis can achieve robust brain activation maps despite task-correlated motion, as seen in Fig 9D. Motion components separate out with similar impact on functional and dynamic images due to motion, as shown in Fig. 9F.

**Figure 9:**
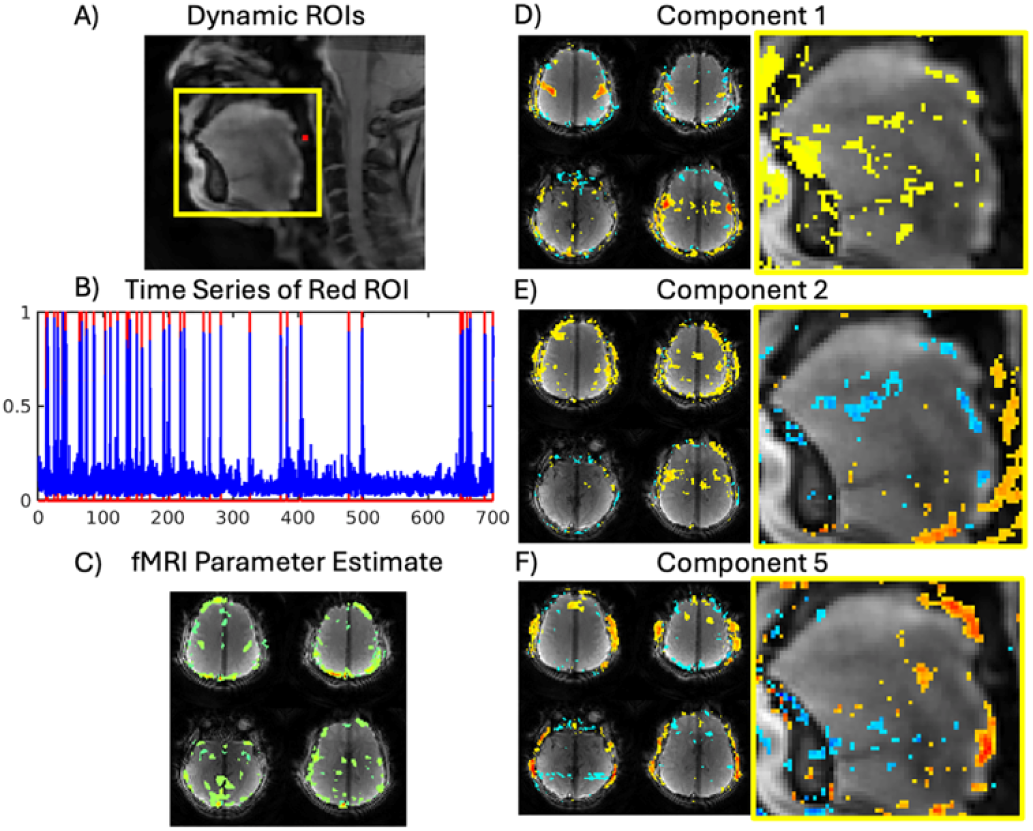
Comparison of standard fMRI and PLSC results in self-paced swallowing task in Subject 4. **A)** Cropped dynamic image frame showing the red pixels as the region of interest for determining swallowing timing for standard fMRI analysis. The yellow box is the ROI used in PLSC analysis. **B)** Time series of the red ROI, blue trace is mean signal and red lines are the detected swallows used in the timing of the task for the standard fMRI analysis. **C)** Parameter estimate from standard fMRI analysis with GLM using swallow timings from slices 8, 16, 22,and 26. **D)-F)** Component 1, 2, and 5 from the PLSC analysis of the fMRI data (all slices) and the dynamic ROI. Components 1 and 2 show functional signals whereas component 5 shows edge voxels for both dynamic and fMRI, indicating motion. Scaling as in Figure 6.

## Discussion

The updated SimulScan sequence achieves 23.75 fps dynamic imaging and a 1.6 s TR functional MRI to provide an unprecedented examination of the central control of swallowing. Importantly, the updated sequence integrates advances in fast dynamic imaging to produce clear images that provide significant information about swallow biomechanics. A strong positive correlation in tracking the 10 key anatomical landmarks over the swallow between two SimulScan runs shows that the dynamic MRI scan can reliably evaluate swallowing kinematics. Additionally, the correlation between VFSS and dynamic images from SimulScan was moderate. Among the landmarks with greater magnitude difference, we believe that mandible, C1, and C4 likely reflect postural differences as the angle of the head laying down in the MRI magnet and during VFSS was slightly different. Although also performed in the supine position, VFSS used a pillow that provided comfort to the subject, as seen in Fig. 3. Differences related to hard palate, hyoid bone, and UES likely reflect differences in annotation strategies as each modality shows these structures differently (61). However, in principle, these differences should not substantially affect within-modality CASM analyses, which characterize the dynamic features of pharyngeal swallowing mechanics.

This analysis shows that the dynamic MRI scan can enable reliable evaluation of the functional anatomy underlying pharyngeal swallow mechanics. Although all subjects in the current study were healthy, this is an important early validation that provides confidence in the future use of this new sequence to evaluate disordered swallows and corresponding brain activations in neurogenic dysphagia. With motion robustness and the use of saliva swallows, this technique has great potential for use with patient populations. SimulScan may enable the characterization of impacts of brain pathology on disruptions to swallowing function in stroke, Parkinson’s Disease, and other clinical conditions, providing unparalleled insights into swallow deficits and plastic responses.

The SimulScan sequence and PLSC analysis routine also showed robustness to motion during the swallow, a challenge that plagues standard fMRI approaches. This is likely due to two aspects of the current analysis: First, since the hemodynamic delay is implemented in aligning dynamic and functional data, immediate motion related to the swallow will not correlate with brain functional activations in the PLSC analysis. We have shifted the dynamic time series so that dynamic motions (which would affect fMRI and dynamic scans at the same time) are no longer aligned between the two time series. Some residual motion correlations are seen appearing mainly in a latter component, but the spatial pattern of correlated pixels, around the complete edges of structures, clearly indicates that these are not neurological components. This approach at separateing motion artifacts is similar to other methods which leverage SVD or independent components to select components in fMRI data that are more related to neural activations instead of artifacts, such as in ICA-AROMA (62). Second, we used a region of interest for the dynamic scans in the PLSC analysis that focused on the oropharyngeal region, ensuring that the dynamic information is related to swallowing activity of interest.

There are additional strategies that could be added to the analysis approach that may improve the results. First, no slice timing correction of the fMRI data in SimulScan was done in the current work. This fits with the strategy for the dynamic data in which we averaged the temporal standard deviation across the entire fMRI TR. Given the short fMRI TR of 1.6 seconds, slice timing is not expected to have a large impact on the functional maps correlated with the swallowing activity. However, this should be evaluated in future analyses, and we will explore leveraging the higher temporal resolution of the dynamic imaging data in the analysis.

Other strategies that will be considered in future work include using the biomechanics extracted from CASM analysis to drive the dynamic information about the swallow in the SimulScan analysis. PLSC methods can incorporate a wide variety of behavioral measures to extract the correlated brain activity. Using behavioral measures more related to functional and disordered swallows, extracted from the high temporal resolution dynamic imaging data, may improve the clinical applicability of the technique.

In conclusion, an updated acquisition and analysis approach for SimulScan has significantly improved the quality and details of information available on the central control of swallowing. With this approach, we have achieved 23.75 fps dynamic imaging simultaneously with functional MRI of the brain at 1.6 s during self-paced saliva swallowing tasks. Using PLSC, correlated components can be extracted from the dynamics of the swallow and the brain activity during this task. This results in a motion-robust analysis of swallowing related activity which has high test-retest reliability, as shown in a repeated scan framework. SimulScan has the potential to unlock studies of central control in swallowing dysfunction in neurogenic dysphagia and guide personalized treatment planning approaches in the future.

## Acknowledgements

Research reported in this publication was supported by the National Institute of Aging (Grant R01AG078513, M-PIs: Sutton and Malandraki). Data acquisition was also supported in part by NIH grant S10 OD012336 (PI: Dydak). The content is solely the responsibility of the authors and does not necessarily represent the official views of the National Institutes of Health. The authors wish to thank all participants, Anna Szlembarski, and the radiation technologists facilitating data collection.

## Notes

### Competing Interest Statement

The authors have declared no competing interest.

